# Improved discrimination of asymmetric and symmetric arginine dimethylation by optimization of the normalized collision energy in LC-MS proteomics

**DOI:** 10.1101/2020.02.14.949255

**Authors:** Nicolas G. Hartel, Christopher Z. Liu, Nicholas A. Graham

**Affiliations:** Mork Family Department of Chemical Engineering and Materials Science, University of Southern California, Los Angeles, CA 90089; Norris Comprehensive Cancer Center, University of Southern California, Los Angeles, CA 90089

## Abstract

Protein arginine methylation regulates diverse biological processes including signaling, metabolism, splicing, and transcription. Despite its important biological roles, arginine methylation remains an understudied post-translational modification. Partly, this is because the two forms of arginine dimethylation, asymmetric dimethylarginine (ADMA) and symmetric dimethylarginine (SDMA), are isobaric and therefore indistinguishable by traditional mass spectrometry techniques. Thus, there exists a need for methods that can differentiate these two modifications. Recently, it has been shown that the ADMA and SDMA can be distinguished by the characteristic neutral loss (NL) of dimethylamine and methylamine, respectively. However, the utility of this method is limited because the vast majority of dimethylarginine peptides do not generate measurable NL ions. Here, we report that increasing the normalized collision energy (NCE) in a higher-energy collisional dissociation (HCD) cell increases the generation of the characteristic NL that distinguish ADMA and SDMA. By analyzing both synthetic and endogenous methyl-peptides, we identify an optimal NCE value that maximizes NL generation and simultaneously improves methyl-peptide identification. Using two orthogonal methyl peptide enrichment strategies, high pH strong cation exchange (SCX) and immunoaffinity purification (IAP), we demonstrate that the optimal NCE increases improves NL-based ADMA and SDMA annotation and methyl peptide identifications by 125% and 17%, respectively, compared to the standard NCE. This simple parameter change will greatly facilitate the identification and annotation of ADMA and SDMA in mass spectrometry-based methyl-proteomics to improve our understanding of how these modifications differentially regulate protein function.

## Introduction

Protein arginine methylation is a post-translational modification (PTM) that regulates the cell cycle^1^, metabolism^2,3^, signal transduction^4–6^, and transcriptional control^7–11^. Arginine methylation is catalyzed by protein arginine methyltransferases (PRMTs) and occurs in three forms, monomethylarginine (MMA), asymmetric dimethylarginine (ADMA), and symmetric dimethylarginine (SDMA). Although all PRMTs can catalyze MMA, ADMA and SDMA are catalyzed by Type I and Type II PRMTs, respectively. Importantly, ADMA and SDMA modifications can differentially regulate biological function. Examples include histone H3, where ADMA and SDMA modification of arginine 2 promotes the formation of heterochromatin and euchromatin, respectively^10,11^.

Despite its important role in human health and disease, arginine methylation remains understudied relative to other PTMs. Because arginine methylation occurs in three forms, analysis of inherently more complicated than with binary PTMs like phosphorylation. An additional obstacle for mass spectrometry (MS)-based methyl-arginine proteomics is that enrichment of methylated proteins and peptides from complex cell lysates is difficult because of the small size and neutral charge of methylation. Finally, MS-based proteomic studies of protein arginine methylation are complicated by the fact that ADMA and SDMA are isobaric and thus indistinguishable via standard MS techniques.

In MS-based proteomics, peptide identification relies on structural information generated by peptide fragmentation using tandem mass spectrometry (MS2). Common modes of fragmentation include collision-induced dissociation (CID), electron-transfer dissociation (ETD), and higher-energy collisional dissociation (HCD). Each method has its biases regarding peptide hydrophobicity, peptide length, and peptide modifications^12,13^. In the popular Q-Exactive series of mass spectrometers, the collision energy of the HCD cell is defined by a single, user-defined value, the normalized collision energy (NCE). The NCE linearly scales the energy applied to each peptide according to the peptide’s precursor mass, enabling the HCD cell to achieve optimum fragmentation across a wide range of parent ion masses^14,15^.

As the collisional energy applied to a peptide increases, the prevalence of ions from multiple fragmentation events also increases^15,16^. Recently, it has been reported that the isobaric PTMs ADMA and SDMA can be distinguished by the characteristic neutral loss (NL) of dimethylamine and methylamine, respectively^17–21^ (Fig. 1A). However, the vast majority of dimethylarginine (DMA) peptides do not generate measurable NL ions, limiting the utility of this method. Here, using synthetic DMA-containing peptides, we show that increasing the NCE improved the frequency of NL generation from DMA. We then tested the effect of NCE on endogenous methyl-peptides and identified the optimal NCE for methyl-peptide identification and NL annotation of ADMA and SDMA. Finally, using two different methyl-peptide enrichment strategies, we demonstrate that the optimal NCE improved ADMA and SDMA assignment by NL by 125% percent while also improving methyl peptide identifications by 17 percent.

**Figure 1.**
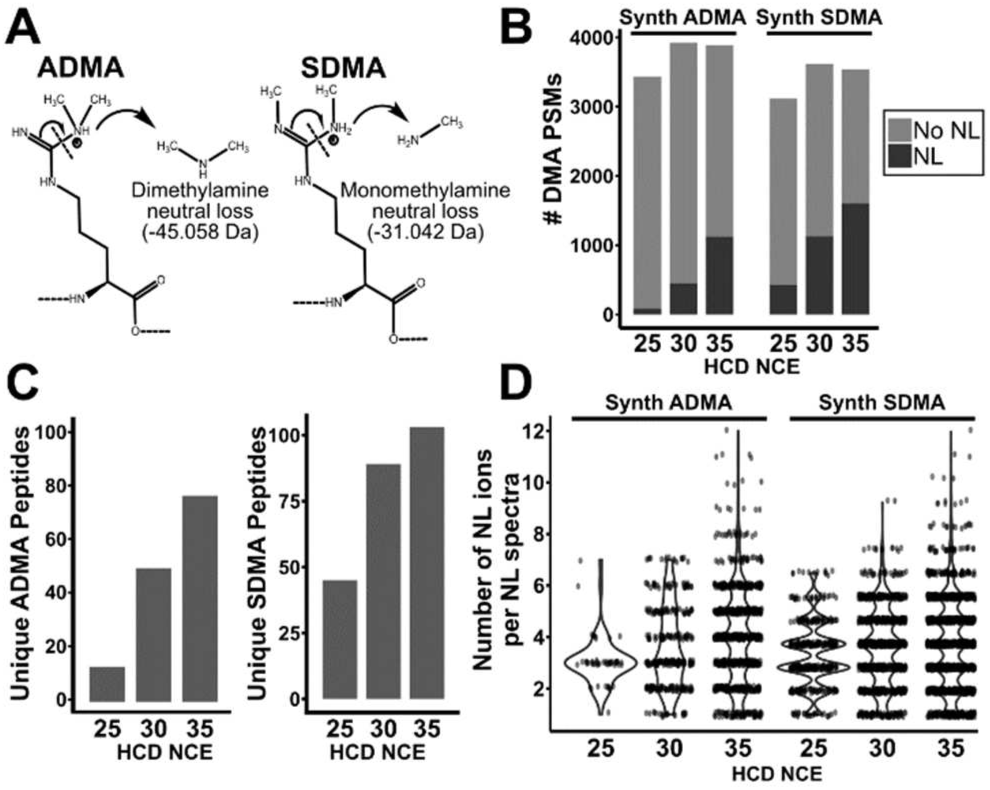
Higher NCE improves generation of NL ions in synthetic ADMA and SDMA peptides. A) Mechanism of neutral loss of dimethylamine and monomethylamine from ADMA and SDMA, respectively. B) Total number of DMA PSMs from a dataset of synthetic ADMA and SDMA peptides. The proportion of PSMs displaying either an ADMA or SDMA neutral loss in the HCD cell of an Orbitrap Fusion Lumos are shown for each NCE. C) Total number of unique peptides with assignable ADMA or SDMA neutral loss for each NCE. D) Violin plot of the number of NL ions present in DMA PSMs for each NCE. Each point represents a PSM with a corresponding number of NL ions observed in that spectra.

## Experimental Section

### Cell Culture

HEK 293T cells were grown in DMEM media (Corning) supplemented with 10% FBS (Omega Scientific) and 100 U/mL penicillin/streptomycin (Thermo Scientific). Cells were cultured at 37 °C in a humidified 5% CO_2_ atmosphere.

### Cell Lysate Preparation

Cells were washed with PBS, scraped, and lysed in 50 mM Tris pH 7.5, 8 M urea, 1 mM activated sodium vanadate, 2.5 mM sodium pyrophosphate, 1 mM β-glycerophosphate, and 100 mM sodium phosphate. Protein concentrations were measured by bicinchoninic acid assay. Lysates were sonicated and cleared by high speed centrifugation and then filtered through a 0.22 μm filter. Proteins were reduced, alkylated, and quenched with 5 mM dithiothreitol, 25 mM iodoacetamide, 10 mM dithiothreitol, respectively. Lysates were four-fold diluted in 100 mM Tris pH 8.0 and digested with trypsin at a 1:100 ratio and then quenched with addition of trifluoroacetic acid to pH 2. Peptides were purified using reverse-phase Sep-Pak C18 cartridges (Waters) and eluted with 30% acetonitrile, 0.1% TFA and then dried by vacuum. Dried peptides were subjected to high pH strong cation exchange or antibody immunoaffinity purification.

### High pH Strong Cation Exchange (SCX)

As described previously^22,23^, 1 mg of digested protein was resuspended in loading buffer (60% acetonitrile, 40% BRUB (5 mM phosphoric acid, 5 mM boric acid, 5 mM acetic acid, pH 2.5) and incubated with high pH SCX beads (Sepax) for 30 minutes, washed with washing buffer (80% acetonitrile, 20% BRUB, pH 9), and eluted into four fractions using elution buffer 1 (60% acetonitrile, 40% BRUB, pH 9), elution buffer 2 (60% acetonitrile, 40% BRUB, pH 10), elution buffer 3 (60% acetonitrile, 40% BRUB, pH 11), and elution buffer 4 (30% acetonitrile, 70% BRUB, pH 12. Eluates were dried, resuspended in 1% trifluoroacetic acid and desalted on STAGE tips^24^ with 2 mg of HLB material (Waters) loaded onto 300 uL tip with a C8 plug (Empore, Sigma).

### Immunoaffinity Purification (IAP)

10 mg of digested proteins were dissolved in 1X immunoprecipitation buffer (50 mM MOPS, 10 mM Na_2_HPO_4_, 50 mM NaCl, pH 7.2, Cell Signaling). Modified asymmetric dimethyl arginine peptides were immunoprecipitated by addition of 40 uL of PTMScan Asymmetric Di-Methyl Arginine Motif Kit (13474, Cell Signaling). Lysates were incubated with PTMScan motif kits for 2 hours at 4 °C on a rotator. Beads were centrifuged and washed two times in 1X immunoprecipitation buffer followed by three washes in water, and modified peptides were eluted with 2 × 50 uL of 0.15% TFA and desalted on STAGE tips with C18 cores (Empore, Sigma). Enriched peptides were resuspended in 50 mM ammonium bicarbonate (Sigma) and subjected to a second digestion with trypsin for 2 hours per the manufacturer’s recommendation, acidified with trifluoroacetic acid to pH 2 and desalted on STAGE tips.

### Mass Spectrometric Analysis

All LC-MS experiments were performed on a nanoscale UHPLC system (EASY-nLC1200, Thermo Scientific) connected to an Q Exactive Plus hybrid quadrupole-Orbitrap mass spectrometer equipped with a nanoelectrospray source (Thermo Scientific). Peptides were separated by a reversed-phase analytical column (PepMap RSLC C18, 2 μm, 100 Å, 75 μm X 25 cm) (Thermo Scientific). For high pH SCX fractions the flow rate was set to 300 nl/min at a gradient starting with 0% buffer B (0.1% FA, 80% acetonitrile) to 29% B in 142 minutes, then washed by 90% B in 10 minutes, and held at 90% B for 3. The maximum pressure was set to 1,180 bar and column temperature was constant at 50 °C. For IAP samples the flow rate was set to 300 nl/min at a gradient starting with 0% buffer B to 25% B in 132 minutes, then washed by 90% B in 10 minutes. Dried SCX fractions were resuspended in buffer A and injected as follows, E1: 1.5 µL/60 µL, E2-4: 5 µL/6 µL. IAP samples were resuspended in 7 µL and 6.5 µL was injected. The effluent from the HPLC was directly electrosprayed into the mass spectrometer. Peptides separated by the column were ionized at 2.0 kV in the positive ion mode. MS1 survey scans for DDA were acquired at resolution of 70k from 350 to 1800 m/z, with maximum injection time of 100 ms and AGC target of 1e6. MS/MS fragmentation of the 10 most abundant ions were analyzed at a resolution of 17.5k, AGC target 5e4, maximum injection time 120 ms for IAP samples, 240 ms for SCX samples, and normalized collision energies of 26, 30, 32, 34, 38, 42, 46, and 50 were tested. Dynamic exclusion was set to 30 s and ions with charge 1 and >6 were excluded. The mass spectrometry proteomics data have been deposited to the ProteomeXchange Consortium (http://proteomecentral.proteomexchange.org) via the PRIDE partner repository with the dataset identifier PXD017193.

### Identification and Quantitation of Peptides

MS/MS fragmentation spectra were searched with Proteome Discoverer SEQUEST (version 2.2, Thermo Scientific) against the in-silico tryptic digested Uniprot *H. sapiens* database with all reviewed with isoforms (release June 2017, 42,140 entries). The maximum missed cleavage rate was set to 5. Trypsin was set to cleave at R and K. Dynamic modifications were set to include mono-methylation of arginine or lysine (R/K, +14.01565), dimethylation of arginine or lysine (R/K, +28.0313), tri-methylation of lysine (K, +42.04695), oxidation on methionine (M, +15.995 Da, and acetylation on protein N-terminus (+42.011 Da). Fixed modification was set to carbamidomethylation on cysteine residues (C, +57.021 Da). The maximum parental mass error was set to 10 ppm and the MS/MS mass tolerance was set to 0.02 Da. Peptides with sequence of seven to fifty amino acids were considered. Methylation site localization was determined by ptm-RS node in Proteome Discoverer, and only sites with localization probability greater or equal to 75% were considered. The False Discovery Rate threshold was set strictly to 0.01 using Percolator node validated by q-value.

### Methyl False Discovery Estimation

The “Decoy PSMs” export from Proteome Discoverer 2.2 was filtered for decoy methyl PSMs and the decoy q-values from the Percolator node were extracted and compared to the target methyl PSM q-values. Target methyl PSMs were removed until a 1% FDR was achieved as described ^25^.

### Neutral Loss Identification in MaxQuant

The modifications SDMA and ADMA were added to MaxQuant’s library with the added mass of dimethyl on arginine and the corresponding neutral loss masses of 31.042 for SDMA and 45.058 for ADMA assigned in the “Neutral Loss” table in Configuration ^17^. The missed cleavage rate was set to 5 and all other settings were kept unchanged. The “FTMS Dependent Losses” setting was checked. All RAW files were searched with monomethyl(K/R), ADMA, SDMA, dimethyl(K), and oxidation of methionine as variable modifications. Carbamidomethylation was kept as a fixed modification. Neutral losses and their masses were extracted from the msms.txt file using an in-house R script. Only target methyl peptides that passed the 1% Methyl FDR filter were considered for analysis. An Andromeda cutoff score of 56 was also used to filter spectra to reduce the number of incorrect assignments. A custom R script was used to remove neutral losses that did not have the corresponding b/y ion present (e.g., if y6* but not y6 was present, the neutral loss was removed). Peptides that displayed SDMA neutral loss with a corresponding MMA site were further checked to see if the neutral loss could be localized to the dimethyl site. If the SDMA neutral loss was not localizable to the dimethyl site, the peptide was removed. A few spectra were confirmed by manual inspection to ensure the accuracy of the Andromeda search. For identified ADMA / SDMA neutral losses, the Andromeda output was matched to Proteome Discoverer data by MS2 scan number.

## Results and Discussion

### Higher NCE improves discrimination of ADMA and SDMA in synthetic peptides

Fragmentation of DMA-containing peptides can result in NL of dimethylamine and monomethylamine from ADMA and SDMA, respectively (Fig. 1A). These NL generate mass shifts of 31.042 Da for ADMA peptides and 45.058 Da for SDMA peptides. To assess the effect of NCE on NL generation, we analyzed a set of 140 synthetic peptides containing either ADMA or SDMA that had been fragmented at 25, 30, or 35 NCE in the HCD cell of an Orbitrap Fusion Lumos (PRIDE ID: PXD009449)^19^. Although the total number of DMA peptide spectral matches (PSMs) was not significantly changed by increasing NCE, the number of PSMs with either ADMA or SDMA NL strongly increased at higher NCE (Fig. 1B). In addition, the total number of unique ADMA and SDMA peptides with NL substantially increased with higher NCE (Fig. 1C). Notably, synthetic SDMA peptides generated more NL than synthetic ADMA peptides. This is potentially because there are two bonds in SDMA whose fragmentation produces the NL, but only one such bond in ADMA. For these synthetic peptides, even NCE 35 did not appear to have reached a maximum in terms of NL generation.

In addition, higher NCE values significantly increased the number of NL ions observed per spectra for both synthetic ADMA and SDMA peptides (Fig. 1D). Comparison of the Andromeda scores for peptides identified at all three NCEs revealed significantly higher scores for spectra collected at higher NCE (Supp. Fig. 1). Increased Andromeda scores at higher NCE likely arise because Andromeda considers the number of NL in calculating the score for each PSM. Taken together, this data supports that increasing NCE improves the ability to distinguish the isobaric isomers ADMA and SDMA by generating the characteristic NL ions without sacrificing peptide sequencing capabilities.

### Optimization of NCE using endogenous DMA peptides

Next, we sought to investigate the effect of higher NCE on NL generation on endogenous DMA peptides enriched from whole cell lysates. We thus enriched methyl-peptides from 293T whole cell lysates using strong-cation exchange chromatography (SCX) at high pH^22,23^. This antibody-free approach enriches methyl peptides based on the increased positive charge of peptides with missed trypsin cleavages at methylarginine. Analyzing the third SCX elution fraction at NCE values ranging from 26 to 42 on a Q-Exactive Plus mass spectrometer (Fig. 2A), we found that the number of DMA PSMs was substantially increased at NCE values of 30 and 34 compared to the standard value of NCE 26 (Fig. 2B and Supp. Table 1). In addition, the number of PSMs with ADMA or SDMA NL greatly increased as the NCE was increased from 26 to 30 and 34. However, at NCEs greater than 34, both the number of identified DMA peptides and the number of PSMs with ADMA or SDMA NL was decreased.

**Figure 2.**
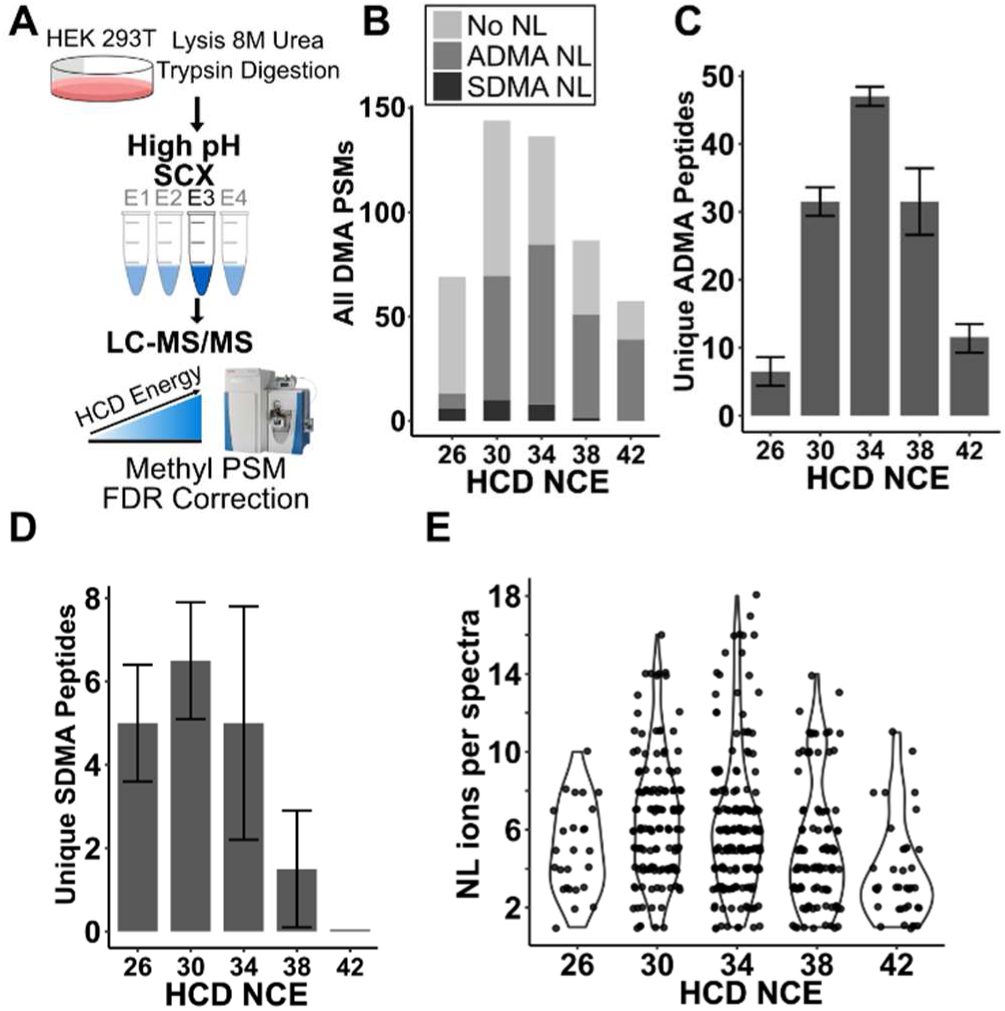
Optimization of NCE for discrimination of ADMA and SDMA in endogenous methyl-peptides. A) Schematic of high pH SCX enrichment of methyl-peptides from 293T cells and analysis of elution fraction three by LC-MS at increasing NCE. B) Total number of DMA PSMs from fraction three of SCX. PSMs with assignable ADMA or SDMA neutral losses are shown. C) Total number of unique ADMA peptides from fraction three of SCX identified at each NCE. D) Total number of unique SDMA peptides from fraction three of SCX identified at each NCE. E) Violin plot of the number of NL ions present in DMA PSMs at each NCE level from fraction three of SCX. Each point represents a PSM with a corresponding number of NL ions within that spectra.

In addition, the number of unique DMA peptides showing ADMA NL increased as the NCE was increased from the standard value of 26 (Fig. 2C). The number of unique DMA peptides with SDMA NL was also slightly increased at NCE 30 compared to the standard NCE value of 26 (Fig. 2C). Notably, the number of peptides with SDMA NL was ∼8 times lower than the number with ADMA NL, even though synthetic SDMA peptides generated more NL than synthetic ADMA peptides (Fig. 1C). This likely reflects the lower abundance of SDMA compared to ADMA *in vivo*^26^. As with synthetic DMA peptides, the number of NL ions per spectra increased up to a maximum at 34 NCE and began diminishing past 34 NCE (Fig. 2D). Lastly, we estimated the methyl false discovery rate (methyl-FDR) for each NCE. The percolator q-value cutoff required to reach a 1% methyl FDR increased as the NCE increased up to 42 NCE (Supp Fig. 2), indicating less methyl decoy spectra were identified at the higher NCEs. Thus, higher NCE can not only improve the chance of NL observation but also reduce the methyl peptide FDR, thereby increasing the total number of identified DMA peptides.

We note the optimum NCE value for the identification and NL generation of endogenous methyl-peptides in this single SCX elution fraction was ∼32. Above NCE 32, the collision energy is likely over-fragmenting peptides, leaving too few ions for peptide identification. In contrast, with synthetic DMA peptides, the number of peptides with observed NL had not saturated even at NCE 35 (Fig. 1). This discrepancy between endogenous and synthetic peptides may occur because of the different MS platforms used (Orbitrap Fusion Lumos v. Q-Exactive Plus) or because of the lower abundance and higher sample complexity of endogenous methyl peptides. Regardless, the fact that previous methyl-proteomics studies used an NCE of 26^22,23^ supports that these studies missed identification and assignment of ADMA/SDMA on the basis of NL for many DMA peptides.

### Optimized NCE improves assignment of ADMA and SDMA spectra in orthogonal methyl peptide enrichment techniques

We next tested whether the optimized NCE value of 32 could improve methyl-peptide identification and ADMA/SDMA assignment in two orthogonal methyl peptide enrichment strategies, high pH SCX and ADMA immunoaffinity purification (IAP). We purified methyl-peptides from 293T cell lysates using both strategies and then analyzed samples at both standard NCE 26 and optimized NCE 32 on a Q-Exactive Plus (Fig. 3A and Supp. Tables 2 and 3). For methyl-peptides purified by high pH SCX, the number of unique DMA peptides identified at NCE 32 increased by ∼20% relative to NCE 26. Additionally, the number of unique DMA peptides with assignable NL increased ∼70% at NCE 32 compared to NCE 26 (Fig. 3B). This increase was primarily driven by identification of ADMA NL.

**Figure 3.**
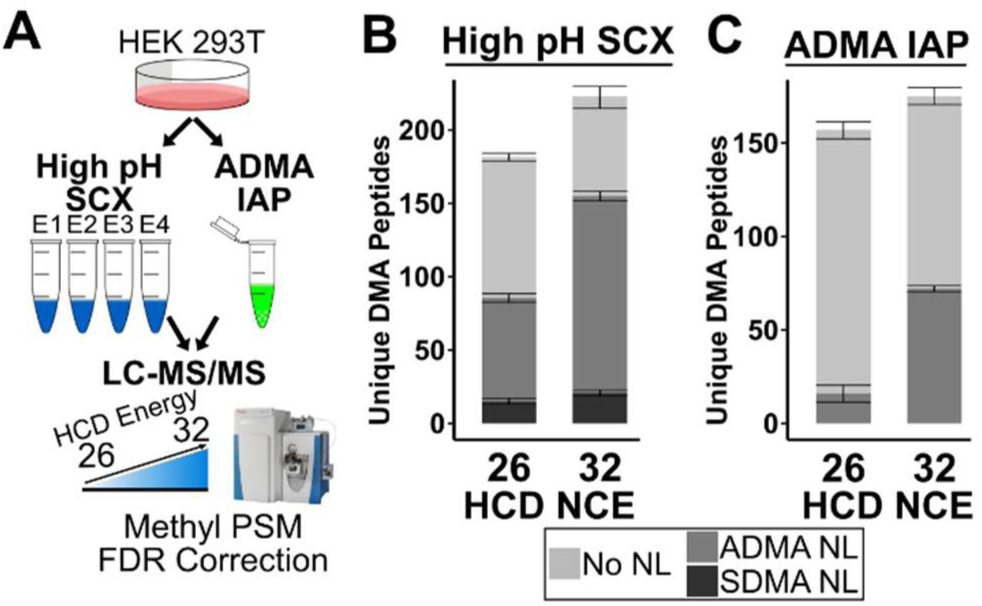
Comparing standard and optimized NCE for high pH SCX and ADMA immunoaffinity methyl-peptide purification. A) 293T cells were lysed, digested, and subject to high pH SCX enrichment and an ADMA immunoaffinity enrichment. Enriched SCX fractions and the ADMA IAP elution were run at a standard NCE of 26 and an ‘optimum’ NCE of 32. B) The number of unique DMA peptides identified across all high pH SCX fractions at 26 and 32 NCE. Peptides with assignable neutral loss are shown. C) The number of unique DMA peptides identified in the ADMA IAP at 26 and 32 NCE. Peptides with assignable neutral loss are shown. One SDMA NL peptide was identified in the ADMA IAP experiment, but this peptide was not included on the chart for clarity.

For methyl-peptides purified by ADMA IAP, increasing the NCE from 26 to 32 did not significantly increase the total number of unique DMA peptides identified. However, increasing the NCE did result in a ∼350% increase of DMA peptides with assignable ADMA NL (Fig. 3C). Only one SDMA NL were observed in these ADMA IAP samples. Notably, there are still ∼25% and ∼50% of peptides in SCX and IAP, respectively, which were not annotated with NL. Thus, there exists further need to optimize methods for discrimination of ADMA and SDMA by MS. Regardless, increasing the NCE from 26 to 32 resulted in increases of ∼125% and ∼17% for NL annotation and methyl-peptide identification.

### Optimized NCE improves confidence of ADMA assignment through higher occurrence of NL across ion series

DMA spectra that contain only a single NL may decrease the reliability of ADMA and SMDA assignment by NL. In addition, false identification of a single NL ion may be increased because NL ions are typically much less abundant than b and y ions in HCD spectra. We thus compared the MS2 spectra of the same DMA peptide collected at 26 and 32 NCE (TAF15 R206, peptide GPMTGSSGGD**<u>R</u>**GGFK where **<u>R</u>** indicates the DMA modified residue). At 26 NCE, GPMTGSSGGD**<u>R</u>**GGFK generated only one identifiable NL (y_11_^+^) (Fig. 4). However, at 32 NCE, the same peptide generated seven identifiable NL ions, all on y ions. As such, the likelihood of correct ADMA assignment for this peptide is highly increased. Furthermore, the generation of multiple NL ions can improve the localization of ADMA/SDMA on peptides that contain multiple arginine residues. This is especially beneficial for methyl-peptides enriched by high pH SCX which frequently contain missed cleavages and multiple methylation sites within the same peptide^23^.

**Figure 4.**
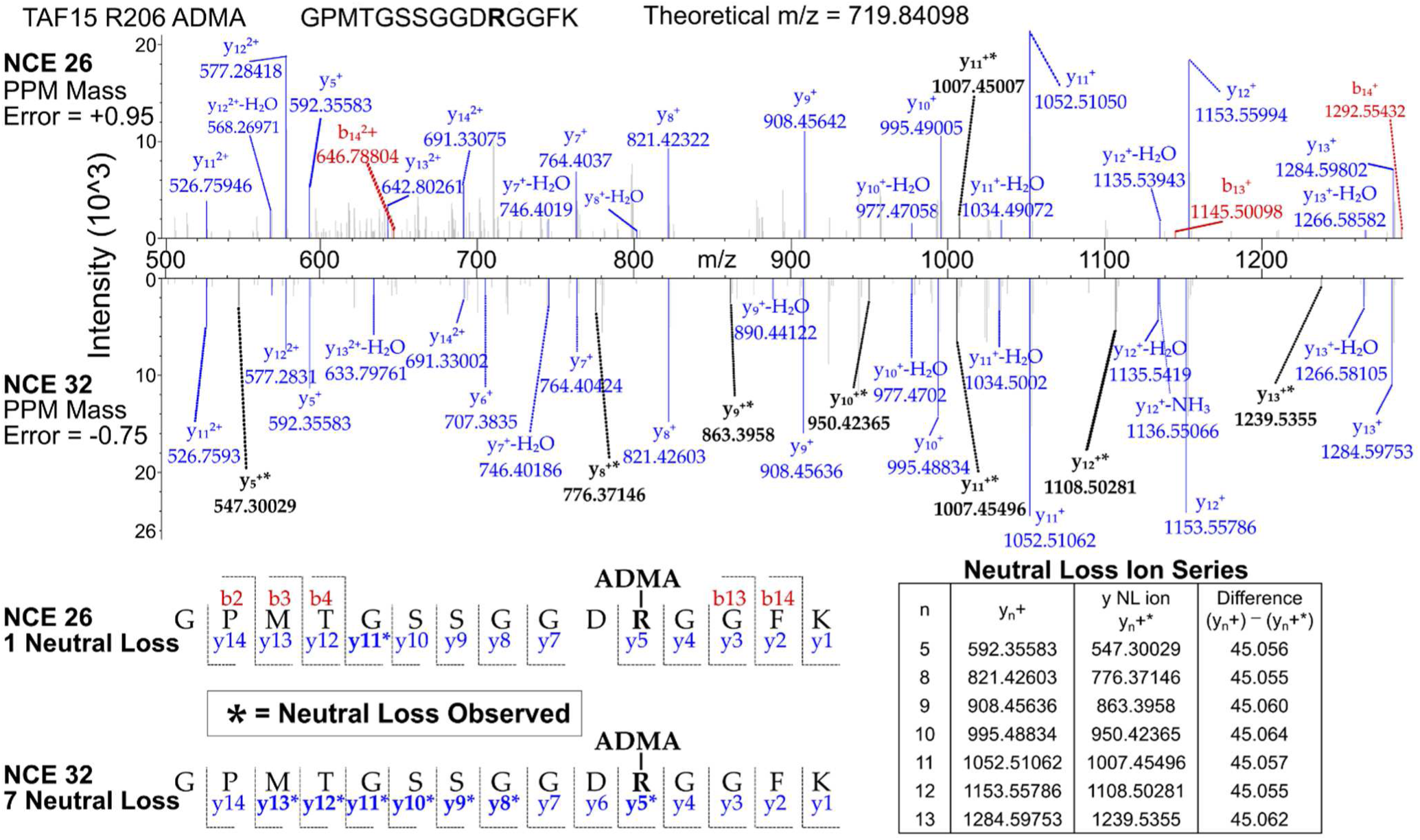
Higher NCE improves occurrence of NL across ion series, improving confidence of ADMA identification. Two MS2 spectra are shown for the same peptide fragmented at 26 NCE (above) and 32 NCE (below). The b and y ions are highlighted in red and blue, respectively, and the NL ions in black. An inset table shows the difference of the mass to charge ratios of the y+ ions and their corresponding NL ions which equal the mass loss of dimethylamine (45.058 Da). The displayed spectra have similar intensities, reducing the effect from higher ion loading in the collision chamber.

## Conclusion

We have demonstrated that using higher NCE during HCD fragmentation improves the assignment of NL for DMA peptides, thereby enabling improved discrimination of the isobaric modifications ADMA and SDMA. In addition, increased NCE improves the total number of DMA peptide identifications in two methyl-peptide enrichment strategies, high pH SCX and IAP. Given the emerging evidence that ADMA and SDMA differentially regulate biological function, this study demonstrates a simple method to improve proteomic studies of protein arginine dimethylation by mass spectrometry.

## AUTHOR INFORMATION

### Author Contributions

NGH and NAG conceived of the project; NGH and CZL conducted the experiments. NGH analyzed data. NGH and NAG wrote the manuscript. All authors have given approval to the final version of the manuscript.

### Notes

The authors declare no competing financial interest.

## ACKNOWLEDGMENT

This work was supported by the USC Viterbi School of Engineering. N.G.H. was partly supported by a Mork Family Departmental Fellowship. C.Z.L. was partly supported by the USC Bridge Undergraduate Science Program.

## REFERENCES

(1) Scoumanne, A.; Zhang, J.; Chen, X. PRMT5 Is Required for Cell-Cycle Progression and P53 Tumor Suppressor Function. Nucleic Acids Res. 2009, 37 (15), 4965–4976. https://doi.org/10.1093/nar/gkp516.

(2) Liu, F.; Ma, F.; Wang, Y.; Hao, L.; Zeng, H.; Jia, C.; Wang, Y.; Liu, P.; Ong, I. M.; Li, B.; et al. PKM2 Methylation by CARM1 Activates Aerobic Glycolysis to Promote Tumorigenesis. Nat. Cell Biol. 2017, cb3630. https://doi.org/10.1038/ncb3630.

(3) Wang, Y.-P.; Zhou, W.; Wang, J.; Huang, X.; Zuo, Y.; Wang, T.-S.; Gao, X.; Xu, Y.-Y.; Zou, S.-W.; Liu, Y.-B.; et al. Arginine Methylation of MDH1 by CARM1 Inhibits Glutamine Metabolism and Suppresses Pancreatic Cancer. Mol. Cell 2016, 64 (4), 673–687. https://doi.org/10.1016/j.mol-cel.2016.09.028.

(4) Xu, J.; Wang, A. H.; Oses-Prieto, J.; Makhijani, K.; Katsuno, Y.; Pei, M.; Yan, L.; Zheng, Y. G.; Burlingame, A.; Brückner, K.; et al. Arginine Methylation Initiates BMP-Induced Smad Signaling. Mol. Cell 2013, 51 (1), 5–19. https://doi.org/10.1016/j.molcel.2013.05.004.

(5) Hsu, J.-M.; Chen, C.-T.; Chou, C.-K.; Kuo, H.-P.; Li, L.-Y.; Lin, C.-Y.; Lee, H.-J.; Wang, Y.-N.; Liu, M.; Liao, H.-W.; et al. Crosstalk between Arg 1175 Methylation and Tyr 1173 Phosphorylation Negatively Modulates EGFR-Mediated ERK Activation. Nat. Cell Biol. 2011, 13 (2), 174–181. https://doi.org/10.1038/ncb2158.

(6) Biggar, K. K.; Li, S. S.-C. Non-Histone Protein Methylation as a Regulator of Cellular Signalling and Function. Nat. Rev. Mol. Cell Biol. 2015, 16 (1), 5–17. https://doi.org/10.1038/nrm3915.

(7) Sylvestersen, K. B.; Horn, H.; Jungmichel, S.; Jensen, L. J.; Nielsen, M. L. Proteomic Analysis of Arginine Methylation Sites in Human Cells Reveals Dynamic Regulation During Transcriptional Arrest. Mol. Cell. Proteomics 2014, 13 (8), 2072–2088. https://doi.org/10.1074/mcp.O113.032748.

(8) Zheng, S.; Moehlenbrink, J.; Lu, Y.-C.; Zalmas, L.-P.; Sagum, C. A.; Carr, S.; McGouran, J. F.; Alexander, L.; Fedorov, O.; Munro, S.; et al. Arginine Methylation-Dependent Reader-Writer Interplay Governs Growth Control by E2F-1. Mol. Cell 2013, 52 (1), 37–51. https://doi.org/10.1016/j.mol-cel.2013.08.039.

(9) Krapivinsky, G.; Krapivinsky, L.; Renthal, N. E.; Santa-Cruz, A.; Manasian, Y.; Clapham, D. E. Histone Phosphorylation by TRPM6’s Cleaved Kinase Attenuates Adjacent Arginine Methylation to Regulate Gene Expression. Proc. Natl. Acad. Sci. 2017, 114 (34), E7092–E7100. https://doi.org/10.1073/pnas.1708427114.

(10) Migliori, V.; Müller, J.; Phalke, S.; Low, D.; Bezzi, M.; Mok, W. C.; Sahu, S. K.; Gunaratne, J.; Capasso, P.; Bassi, C.; et al. Symmetric Dimethylation of H3R2 Is a Newly Identified Histone Mark That Supports Euchromatin Maintenance. Nat. Struct. Mol. Biol. 2012, 19 (2), 136–144. https://doi.org/10.1038/nsmb.2209.

(11) Kirmizis, A.; Santos-Rosa, H.; Penkett, C. J.; Singer, M. A.; Vermeulen, M.; Mann, M.; Bähler, J.; Green, R. D.; Kouzarides, T. Arginine Methylation at Histone H3R2 Controls Deposition of H3K4 Trimethylation. Nature 2007, 449 (7164), 928–932. https://doi.org/10.1038/nature06160.

(12) Huang, T.-Y.; McLuckey, S. A. Gas-Phase Chemistry of Multiply Charged Bioions in Analytical Mass Spectrometry. Annu. Rev. Anal. Chem. Palo Alto Calif 2010, 3, 365–385. https://doi.org/10.1146/annurev.anchem.111808.073725.

(13) Elviri, L. 7 ETD and ECD Mass Spectrometry Fragmentation for the Characterization of Protein Post Translational Modifications; 2012.

(14) Michalski, A.; Damoc, E.; Hauschild, J.-P.; Lange, O.; Wieghaus, A.; Makarov, A.; Nagaraj, N.; Cox, J.; Mann, M.; Horning, S. Mass Spectrometry-Based Proteomics Using Q Exactive, a High-Performance Benchtop Quadrupole Orbitrap Mass Spectrometer. Mol. Cell. Proteomics 2011, 10 (9). https://doi.org/10.1074/mcp.M111.011015.

(15) Zhang, Y.; Ficarro, S. B.; Li, S.; Marto, J. A. Optimized Orbitrap HCD for Quantitative Analysis of Phosphopeptides. J. Am. Soc. Mass Spectrom. 2009, 20 (8), 1425–1434. https://doi.org/10.1016/j.jasms.2009.03.019.

(16) Olsen, J. V.; Macek, B.; Lange, O.; Makarov, A.; Horning, S.; Mann, M. Higher-Energy C-Trap Dissociation for Peptide Modification Analysis. Nat. Methods 2007, 4 (9), 709–712. https://doi.org/10.1038/nmeth1060.

(17) Musiani, D.; Bok, J.; Massignani, E.; Wu, L.; Tabaglio, T.; Ippolito, M. R.; Cuomo, A.; Ozbek, U.; Zorgati, H.; Ghoshdastider, U.; et al. Proteomics Profiling of Arginine Methylation Defines PRMT5 Substrate Specificity. Sci Signal 2019, 12 (575), eaat8388. https://doi.org/10.1126/scisignal.aat8388.

(18) Nabity, M. B.; Lees, G. E.; Boggess, M. M.; Yerramilli, M.; Obare, E.; Yerramilli, M.; Rakitin, A.; Aguiar, J.; Relford, R. Symmetric Dimethylarginine Assay Validation, Stability, and Evaluation as a Marker for the Early Detection of Chronic Kidney Disease in Dogs. J. Vet. Intern. Med. 2015, 29 (4), 1036–1044. https://doi.org/10.1111/jvim.12835.

(19) Zolg, D. P.; Wilhelm, M.; Schmidt, T.; Medard, G.; Zerweck, J.; Knaute, T.; Wenschuh, H.; Reimer, U.; Schnatbaum, K.; Kuster, B. ProteomeTools: Systematic Characterization of 21 Post-Translational Protein Modifications by LC-MS/MS Using Synthetic Peptides. Mol. Cell. Proteomics 2018, mcp.TIR118.000783. https://doi.org/10.1074/mcp.TIR118.000783.

(20) Brame, C. J.; Moran, M. F.; McBroom-Cerajewski, L. D. B. A Mass Spectrometry Based Method for Distinguishing between Symmetrically and Asymmetrically Dimethylated Arginine Residues. Rapid Commun. Mass Spectrom. 2004, 18 (8), 877–881. https://doi.org/10.1002/rcm.1421.

(21) Rappsilber, J.; Friesen, W. J.; Paushkin, S.; Dreyfuss, G.; Mann, M. Detection of Arginine Dimethylated Peptides by Parallel Precursor Ion Scanning Mass Spectrometry in Positive Ion Mode. Anal. Chem. 2003, 75 (13), 3107–3114.

(22) Wang, K.; Dong, M.; Mao, J.; Wang, Y.; Jin, Y.; Ye, M.; Zou, H. Antibody-Free Approach for the Global Analysis of Protein Methylation. Anal. Chem. 2016, 88 (23), 11319–11327. https://doi.org/10.1021/acs.analchem.6b02872.

(23) Hartel, N. G.; Chew, B.; Qin, J.; Xu, J.; Graham, N. A. Deep Protein Methylation Profiling by Combined Chemical and Immunoaffinity Approaches Reveals Novel PRMT1 Targets. Mol. Cell. Proteomics 2019, 18 (11), 2149–2164. https://doi.org/10.1074/mcp.RA119.001625.

(24) Rappsilber, J.; Ishihama, Y.; Mann, M. Stop and Go Extraction Tips for Matrix-Assisted Laser Desorption/Ionization, Nanoelectrospray, and LC/MS Sample Pretreatment in Proteomics. Anal. Chem. 2003, 75 (3), 663–670. https://doi.org/10.1021/ac026117i.

(25) Elias, J. E.; Gygi, S. P. Target-Decoy Search Strategy for Mass Spectrometry-Based Proteomics. Methods Mol. Biol. Clifton NJ 2010, 604, 55–71. https://doi.org/10.1007/978-1-60761-444-9_5.

(26) Dhar, S.; Vemulapalli, V.; Patananan, A. N.; Huang, G. L.; Di Lorenzo, A.; Richard, S.; Comb, M. J.; Guo, A.; Clarke, S. G.; Bedford, M. T. Loss of the Major Type I Arginine Methyl-transferase PRMT1 Causes Substrate Scavenging by Other PRMTs. Sci. Rep. 2013, 3. https://doi.org/10.1038/srep01311.

