## Supplemental Figures for "Improved discrimination of asymmetric and symmetric arginine dimethylation by optimization of the normalized collision energy in LC-MS proteomics"

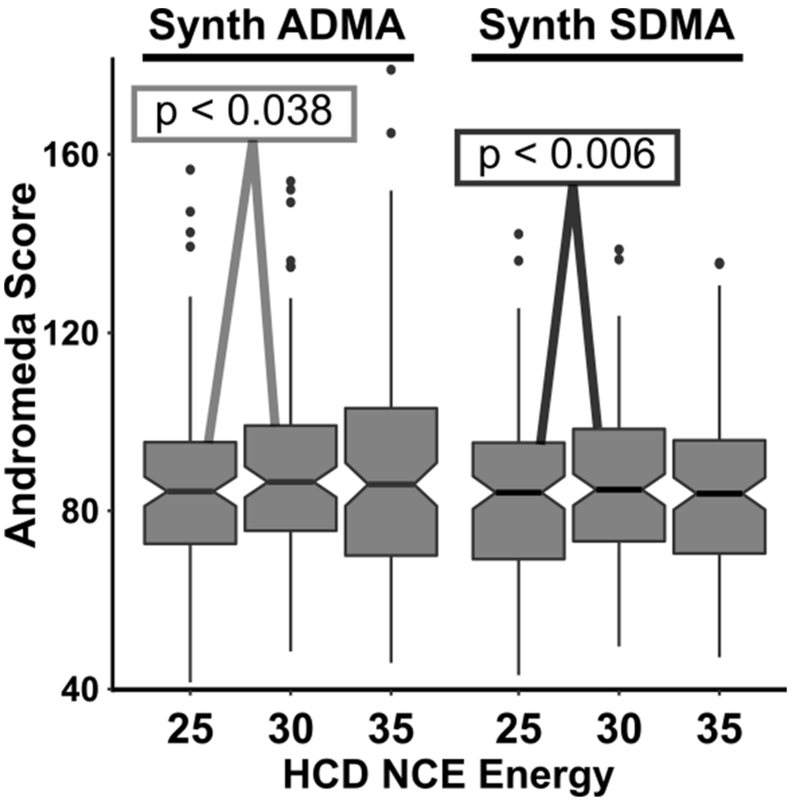


**Supp. Fig. 1. Higher NCE improves Andromeda scores of synthetic ADMA and SDMA peptides.** Boxplots of the Andromeda scores of synthetic DMA peptides fragmented at each NCE. The difference in scores at 25 and 30 NCE were significant by Student's paired t-test ($p < 0.038$ for ADMA, $p < 0.006$ for SDMA). Here, Andromeda was configured to consider NL as part of its scoring algorithm (see Experimental Section).

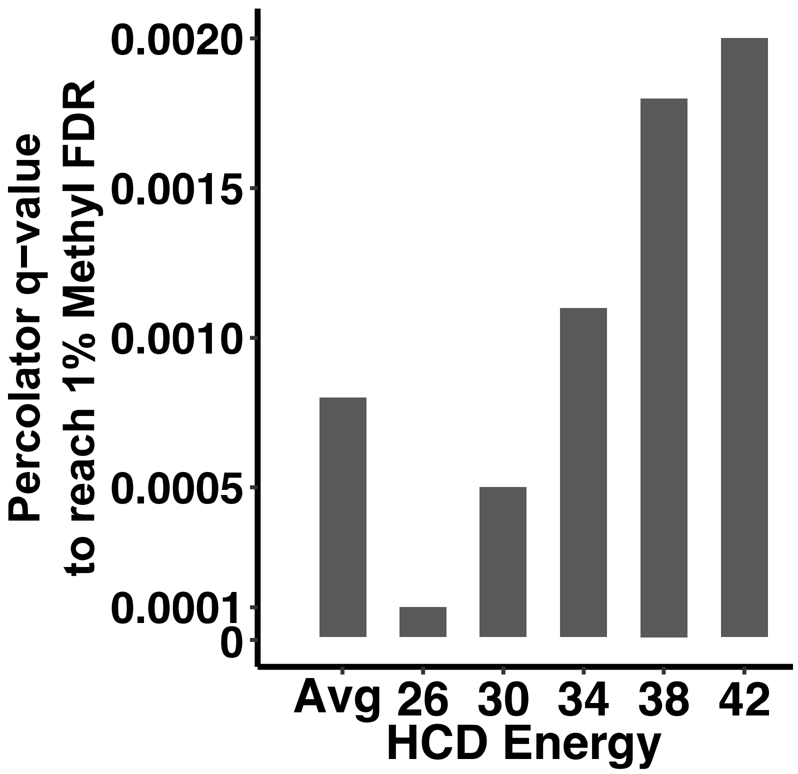


**Supp. Fig. 2. Higher NCE improves methyl PSM false discovery rate by increasing the percolator q-value needed to reach a 1% methyl FDR**

Barchart of the q-value from the Percolator node in Proteome Discoverer 2.2 needed to attain a 1% methyl FDR for each energy. The q-values of the decoy PSMs from the "Decoy PSMs" tab of Proteome Discoverer were extracted and used to estimate the methyl FDR from the q-values in the target dataset. Higher NCEs reduced the number of methyl decoys present, leading to improved q-values for the 1% methyl FDR cutoff. For reference a percolator q-value of 0.01 represents a 1% peptide FDR which is not strict enough for methyl peptides which suffer from higher rates of FDR (1)

### Supplemental References

1. Hart-Smith, G., Yagoub, D., Tay, A. P., Pickford, R., and Wilkins, M. R. (2016) Large Scale Mass Spectrometry-based Identifications of Enzyme-mediated Protein Methylation Are Subject to High False Discovery Rates. *Mol. Cell. Proteomics*. **15**, 989–1006
